# Therapeutic activation of virus-specific resident memory T cells within the glioblastoma microenvironment

**DOI:** 10.1101/2021.09.06.458939

**Authors:** Jianfang Ning, Noah V. Gavil, Shaoping Wu, Sathi Wijeyesinghe, Eyob Weyu, Jun Ma, Ming Li, Florina-Nicoleta Grigore, Sanjay Dhawan, Alexander G. J. Skorput, Shawn C. Musial, Clark C. Chen, David Masopust, Pamela C. Rosato

## Abstract

Glioblastoma multiforme (GBM) is among the most aggressive, treatment resistant cancers, and despite standard of care surgery, radiation and chemotherapy, is invariably fatal. GBM is marked by local and systemic immunosuppression, contributing to resistance to existing immunotherapies that have had success in other tumor types. Memory T cells specific for previous infections reside in tissues throughout the host and these tissue resident memory T cells (T_RM_) are capable of rapid and potent immune activation. Here, we show that virus-specific memory CD8+ T cells expressing tissue resident markers populate the mouse and human glioblastoma microenvironment. Reactivating virus-specific T_RM_ through intra-tumoral delivery of adjuvant-free virus-derived peptide triggered local immune activation. This delivery translated to anti-neoplastic effects, which improved survival in a murine glioblastoma model. Our results indicate that virus-specific T_RM_ are a significant part of the glioblastoma immune microenvironment and can be leveraged to promote anti-tumoral immunity.

## Introduction

Glioblastoma multiforme (GBM) is a lethal form of malignant brain tumor and remains refractory to immunotherapies that have transformed the treatment of several cancers. With current standard of care, surgery followed by radiation, chemotherapy, and electric field therapy, median patient survival is less than 18 months. Glioblastoma intra-tumorigenicity and associated immunosuppression within the tumor microenvironment presents unique challenges for therapeutic development (1,2). The presence of the blood-brain and blood-tumor barrier that restrict penetration of large macromolecules further limits antibody-based immunotherapies.

We recently described a novel form of immunotherapy that builds on the activation of virus-specific memory T cells abundant in tumors with no known viral etiology (3–8). This therapy stems from the potent capacity of tissue resident memory T cells (T_RM_) to execute a ‘sensing and alarm’ function upon antigen re-exposure (9). T_RM_ are a subset of memory T cells that reside within tissues, locally patrolling for reinfection and rarely reentering circulation (10,11). Once reactivated, T_RM_ produce pro-inflammatory cytokines that trigger local immune stimulation, and chemokines that recruit innate and adaptive immunity to the site of reactivation (9,12,13). This T_RM_ ‘sensing and alarm’ function can be triggered in mouse models of melanoma through delivery of viral-derived peptides alone. This form of immunotherapy is termed peptide alarm therapy (PAT) and leads to a significant reduction of tumor growth. When PAT is combined with PD-L1 checkpoint blockade, melanoma tumor burden can be entirely eliminated (3). It remains unclear if this strategy can translate to tumors where the immunosuppressive environment dominates, such as GBM.

In this study, we sought to assess the abundance and phenotype of virus-specific T_RM_ in mouse and human GBM tumors with the goal of harnessing their functions as a GBM tumor immunotherapy. Memory T cells specific for influenza A virus (IAV), Epstein-Barr virus (EBV), and/or cytomegalovirus (CMV) were present in all clinical glioblastoma samples studied. These T cells expressed phenotypic markers of tissue residency (CD69 and CD103) and were able to respond to viral peptide, triggering immune activation in explanted patient tumors. Virus-specific T_RM_ were also present in the microenvironment of murine glioblastoma models. Reactivation of these T_RM_ via PAT induced immune activation, with associated improved survival. These findings establish virus-specific T_RM_ as a part of the glioblastoma tumor immune environment and provide a foundation for harnessing them as a tumor immunotherapy.

## Results

### Virus-specific CD8+ T cells populate human GBM tumors

To determine the extent to which virus-specific CD8+ T cells occupy human glioblastomas, we constructed HLA-A^*^02:01 specific tetramers loaded with immunodominant peptides derived from the common viral infections influenza A virus (IAV), cytomegalovirus (CMV) and Epstein-Barr virus (EBV). We stained T cells isolated from patient tumors following standard of care surgical resection. Memory CD8+ T cells specific for common viral infections consistently populated glioblastoma tumors and often, T cells specific for a single viral epitope made up over 1% of the total CD8+ T cell population in the tumor (Figure 1A, 1B). Of note, in all patients where a population of virus-specific T cells was identified in blood (when available), a corresponding population was identified in tumor samples when sufficient tissue was provided for analysis. We found that most antiviral T cells, regardless of viral specificity, expressed CD69 and a subset also expressed CD103 (Figure 1C, 1D). CD69 and CD103 are canonical T_RM_ markers and were not expressed by T cells in venous blood, suggesting that these cells were within tumors and not blood contaminants (14). Using in situ tetramer staining, we visually confirmed that virus-specific T cells were indeed in the tissue, as illustrated by localization outside of CD31+ vascular endothelial cells (Figure 1E). These data demonstrate the presence of virus-specific T_RM_ in human GBM tumors.

**Figure 1.**
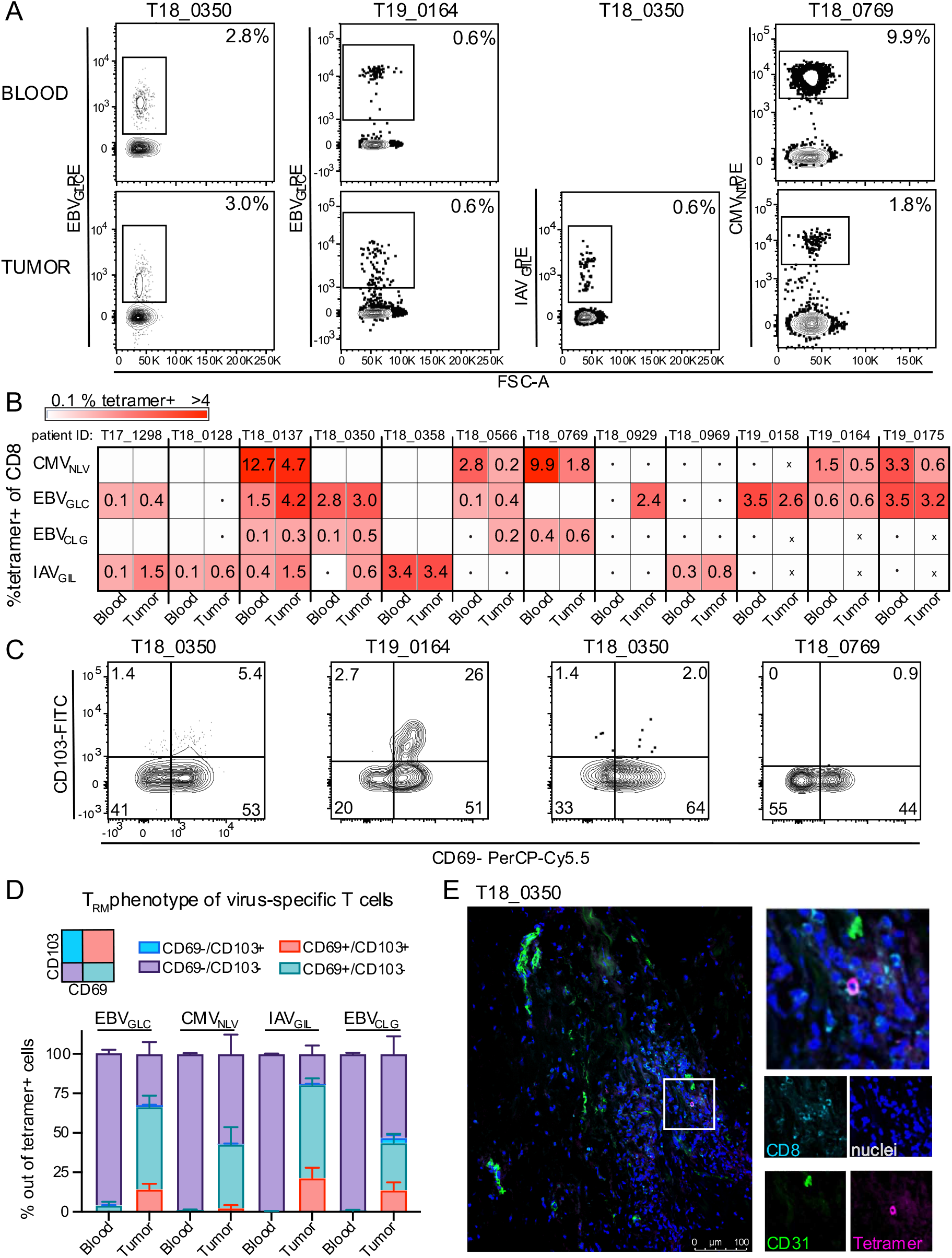
Virus-specific memory CD8+ T cells populate GBM and have a T_RM_ phenotype. **A)** Patient glioblastoma tumors and paired blood stained for HLA-A^*^02+ tetramers specific for EBV (EBV_GLC_ and EBV_CLG_), CMV (CMV_NLV_), and Influenza A virus (IAV_GIL_). Gated on CD8+CD3+ cells. **B)** The frequency (%) of tetramer-positive populations out of CD8+/CD3+ T cells in paired blood and glioblastoma tumors across multiple patients. Dots indicate not enough total CD8+ T cells were acquired to get an accurate measurement, X indicates data was not collected; Blank indicates tetramer+ cells were not detected. **C)** Phenotype of tetramer+ cells gated on in **(A). D)** quantification of CD69/CD103 phenotype of tetramer+ cells in all patient populations in blood and paired tumors. **E)** Representative in situ tetramer immunofluorescence staining of glioblastoma tumor. Magenta, EBV/Flu tetramer; Teal, CD8; Blue, 4’,6-diamidino-2-phenylindole (DAPI)-stained nuclei; Green, CD31. White scale bar=100μm.

### Virus-specific T_RM_ perform sensing and alarm functions in human GBM

T_RM_ have been described as having a “sensing and alarm” function; upon antigen recognition in tissues, they trigger a broad local antimicrobial response (9,12,13). The GBM tumor environment is notoriously immunosuppressive and, given the T_RM_ -phenotype of virus-specific T cells in GBM tumors, we tested if these cells could reactivate and perform sensing and alarm functions to reverse this suppression. We first established ex vivo organotypic slice cultures with five HLA-A2+ GBM tumors. This method preserves the tumor environment while promoting tissue survival through culturing on transwell inserts, allowing for sufficient oxygenation (15). Each patient was screened for tetramer-positive cells in blood and based on these results, we added relevant viral peptides or vehicle control to autologous slice cultures (Figure 2A). By adding free peptide comprised only of CD8+ T cell epitopes, we obviated the need for antigen presenting cell processing and were able to attribute any transcriptional changes observed specifically to antiviral CD8+ T cell activation within the tumor. Nine hours following peptide addition, we harvested tumor slices and performed RNAseq on the whole tissue. We found significant differences in gene expression between control and viral peptide treatment in 4 of the 5 tumors (Figure 2B). Several upregulated genes were the same as previously published in healthy tissue upon T_RM_ reactivation including IFNγ, and the chemokines, CXCL9, CXCL10 and CXCL11 (Figure 2B) (12,13). We performed Ingenuity Pathway Analysis on the gene set with the most differentially expressed genes (T18_0969) and identified STAT1 regulated interferon signaling and antiviral pathways (Figure 2C). Further analysis revealed an upregulation of functions and pathways important for antiviral responses and lymphocyte migration (Figure 2D and E). We also observed signatures such as “Axonal Guidance Signaling” and “Synaptogenesis Signaling Pathway”, reflective of the tissue of origin. In all, these data indicate that virus-specific T_RM_ in human GBM tumors can perform sensing and alarm functions and thus may be a tractable therapeutic target for transforming the suppressive tumor environment.

**Figure 2.**
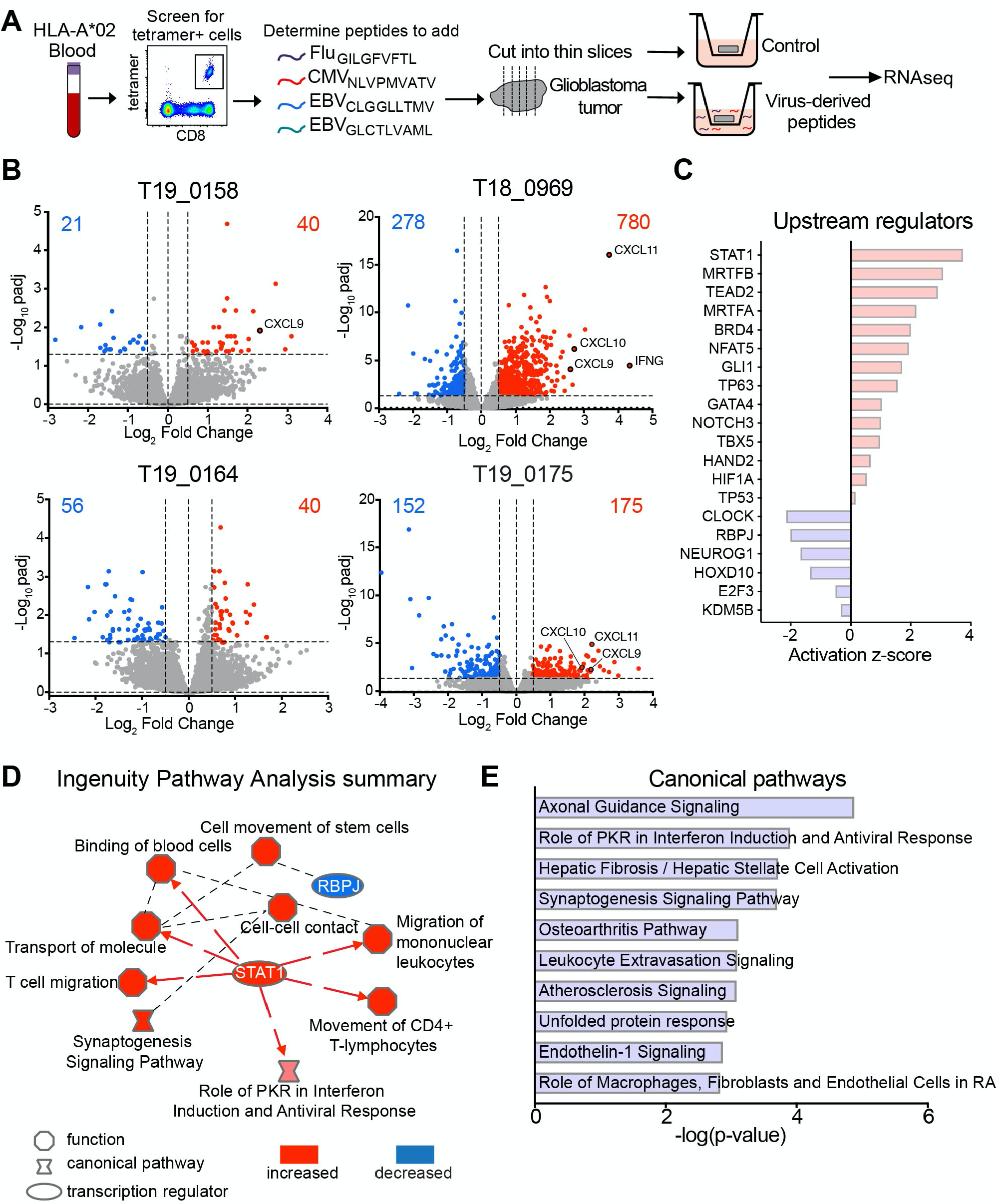
Virus-specific T_RM_ perform sensing and alarm functions in human GBM. **A**) Schematic of experimental set-up. **B)** Volcano plot of 4 patient tumors showing differentially expressed genes in viral peptide treated tumors versus control. **C-E)** Ingenuity Pathway Analysis results from patient T18_0969; **C)** Upstream regulators **D)** Graphical summary of IPA results: Blue: inhibited node z-score<=2, Red: activated node z-score >=2 **E)** Upregulated Canonical Pathways.

### Establishment of virus-specific T_RM_ in mouse GBM following diverse infections

We hypothesized that reactivating T_RM_ with viral peptides *in vivo* could trigger immune activating alarm functions and promote anti-tumor responses. To test this, we first sought to establish a mouse model of GBM which harbors virus-specific memory T cells, mimicking what we observed in humans. Because of the nature of typical pre-clinical mouse models that are housed in specific pathogen free conditions, thus lacking physiologic infectious experience, there are very few memory T cells established in nonlymphoid tissues (16). To establish a trackable population of virus-specific memory T cells, we took advantage of transgenic T cells specific for model antigens. We transferred Thy1.1 or CD45.1 naïve transgenic CD8+ T cells specific for the ovalbumin protein (ova), termed OT-I T cells, or the gp33 epitope of lymphocytic choriomeningitis virus (LCMV), termed P14 T cells, into CD45.2 congenically distinct wild type C57BL/6 host mice. One day after T cell transfer, we infected mice with pathogens of interest expressing the antigens ova or gp33; LCMV, Influenza A virus strain PR8 expressing gp33 (PR8_gp33_), or vesicular stomatitis virus (VSV_ova_) intranasally, or VSV_ova_ intravenously (Figure 3). These infections result in the durable establishment of broadly distributed memory T cells, including T_RM_ in the brain (17–21). Greater than 30 days post-infection (referred to as immune memory mice), we then established GBM tumors using an orthotopic model by intracranially implanting syngeneic GBM cell lines. GL261 is a commonly used GBM cell line which harbors a KRAS mutation, p53 mutations and high expression of c-myc (22). GL261 cells were implanted intracranially into immune memory mice and we assessed T cell infiltration by histology at 14-16 days post-GL261 implantation. Irrespective of the virus or route of infection, we found abundant virus-specific OT-I and P14 T cells in GBM tumors (Figure 3A-D). This observation was also seen in a second tumor model using the 005 cell line, which has HRAS and AKT mutations, and was recently shown to more closely resemble the immune landscape of human glioblastoma (Figure 3D) (23,24). In both the GL261 and 005 models, we observed virus-specific memory T cells expressing CD69 and a subset co-expressed CD69 and CD103 (Figure 3E, F). This T_RM_ phenotype was most prominent in IAV and VSV intranasal infections, but all infection modalities produced T_RM_ that mimic what we observed in human GBM tumors (Figure 3E, F). In summary, memory CD8+ T cells established by diverse viral infections populate mouse GBM models and can express markers associated with tissue residency.

**Figure 3.**
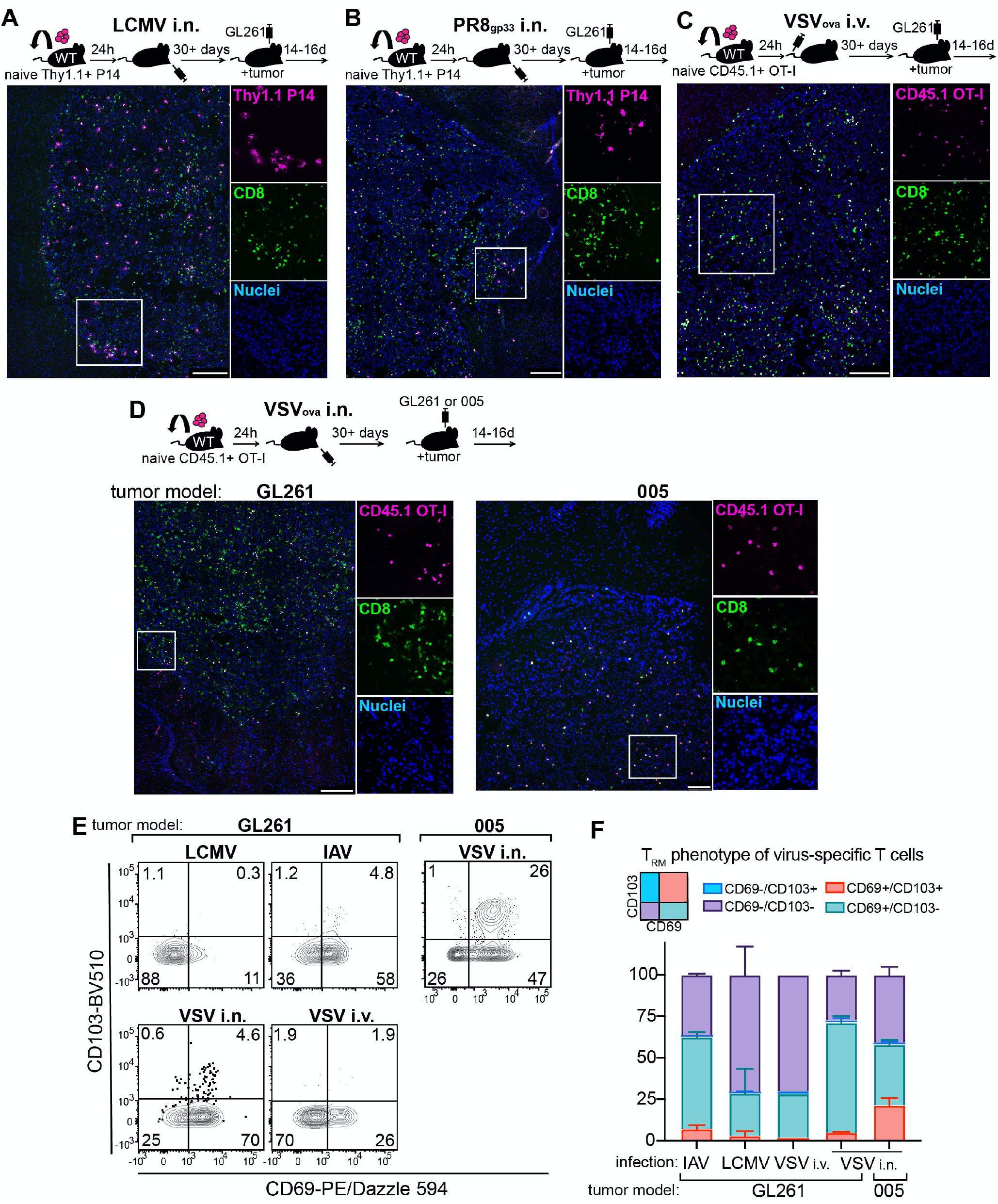
Establishment of T_RM_ in mouse GBM following diverse infections. **A-D** Schematic of experimental set-up utilizing transgenic T cells specific for antigens expressed by concurrent indicated infection, with corresponding immunofluorescence staining of GBM tumor. Magenta, transgenic T cells (P14 or OT-I); Green, CD8; Blue, DAPI stained nuclei. Scale bars=100um. **E)** Representative flow plots gated on transgenic T cells within GBM tumors. **F)** Quantification of E indicating % of total P14 or OT-I populations expressing CD69/CD103.

### Virus-specific T_RM_ can perform sensing and alarm functions in mouse GBM

We next tested if virus-specific T cells can be reactivated in response to cognate viral peptides in the highly immunosuppressive GBM tumor environment. We focused on the intranasal VSV_ova_ infection model as it generated antiviral T_RM_ in tumors that more closely resembled those found in humans. We injected viral (SIINFEKL) or irrelevant control (gp33) peptides intracranially into GL261 or 005 tumors and assessed T_RM_ activation nine hours later (Figure 4A). Consistent with T_RM_ activation, we found that OT-I T cells upregulated IFNγ and CD25 (IL-2Ra) in both tumor models, although a greater percentage of cells appeared activated in GL261 than in 005 tumors (Figure 4B, E). We assessed OT-I activation at a later timepoint, 48 hours post-peptide injection, and found an upregulation of cytotoxic granzyme B and the proliferation marker, Ki67, in GL261 tumors which was not seen in 005 tumors at this timepoint (Figure 4C, F). Despite the increase in Ki67, however, we did not observe an increase in the number of virus-specific T_RM_ 48 hours later (Figure 4D, G).

**Figure 4.**
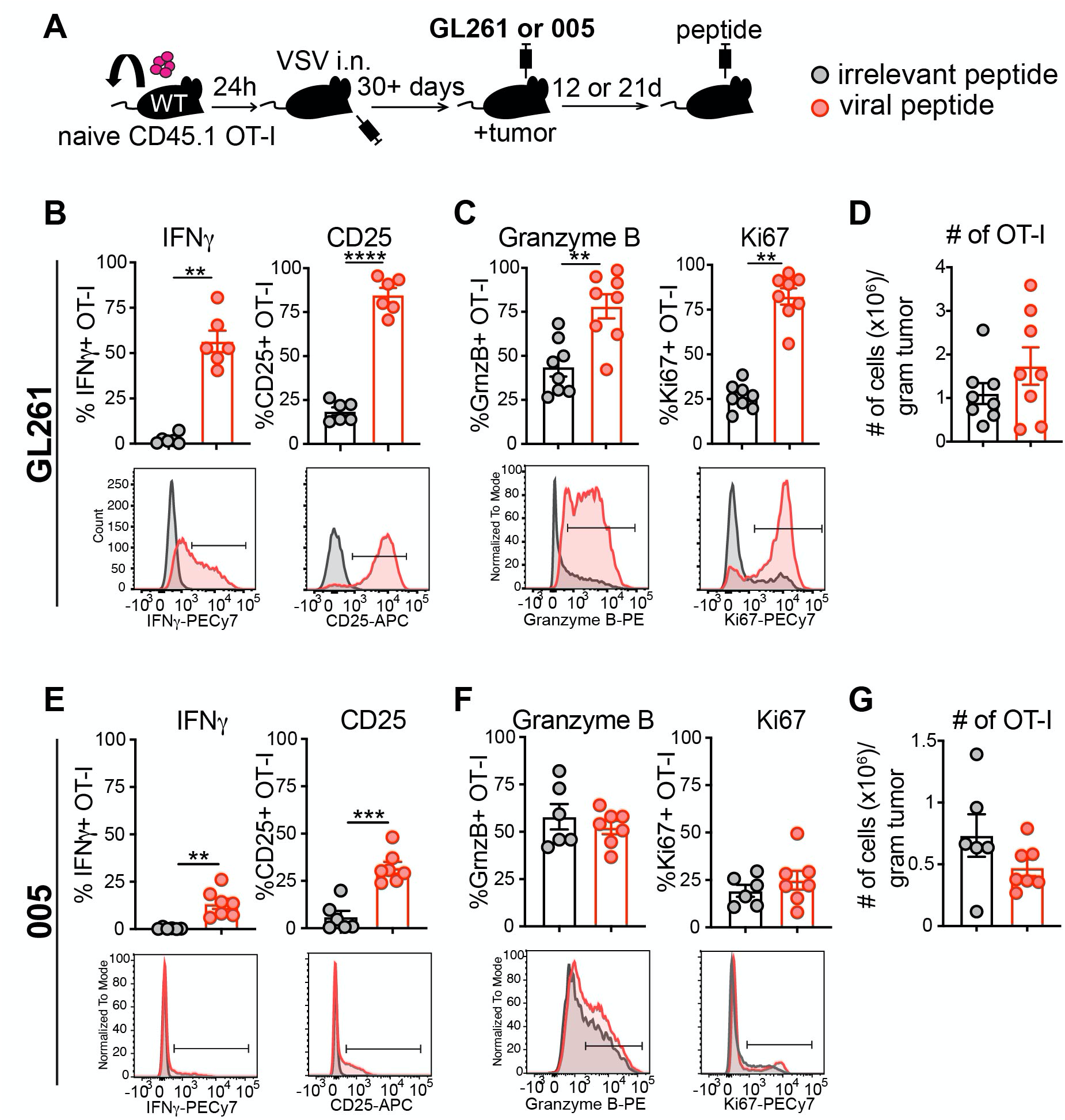
Virus-specific T_RM_ reactivate in mouse GBM. **A**) Schematic of experimental set-up. **B, E)** Proportion of IFNγ+, and CD25+ OT-I in GL261 (B) or 005 (E) tumors 9h following intratumoral injection of irrelevant (grey) or viral SIINFEKL peptide (red). Representative flow plots bellow graphs. **C, F)** Proportion of OT-I T cells expressing granzyme B and Ki67 48 hours post peptide injection of GL261 (C) or 005 (F) tumors, with representative flow plots bellow. **D)** Quantification by flow cytometry of OT-I T cells in GL261 (D) or 005 (G) tumors 48h post-challenge with control, or viral SIINFEKL peptide. Statistical significance was determined by unpaired two-tailed t-test (**B**, CD25 and **C**) and unpaired two-tailed Mann-Whitney test (**B**, IFNγ) where ^**^p<0.01, ^***^p<0.001, ^****^p<0.0001.

We next assessed if T_RM_ reactivation triggered activation of surrounding immune cells, as previously observed in healthy murine skin and female reproductive tract, and in autochthonous and orthotopic mouse models of melanoma (3,12,13). We observed that in both GL261 and 005 tumor models, as a consequence of antigen recognition by CD8+ T_RM_, NK cells became activated, upregulating granzyme B within 48h (Figure 5A). This correlated with a modest increase in Ki67, but paradoxically, a decrease in NK cell numbers in GL261 tumors with no difference in 005 tumors (Figure 5A). We assessed activation of non-OT-I “bystander” memory CD8+ T cells and found an increase in granzyme B expression in T cells isolated from GL261 tumors, but not 005, and no changes in Ki67 or total numbers of cells (Figure 5B). Like NK cells, we observed a small but significant decrease in non-OT-I CD8+ T cell numbers in GL261 tumors (Figure 5B). These data demonstrate that virus-specific T_RM_ can be reactivated in GBM tumors, and this can lead to immune activation of other cell types, albeit more so in the GL261 model than 005.

**Figure 5.**
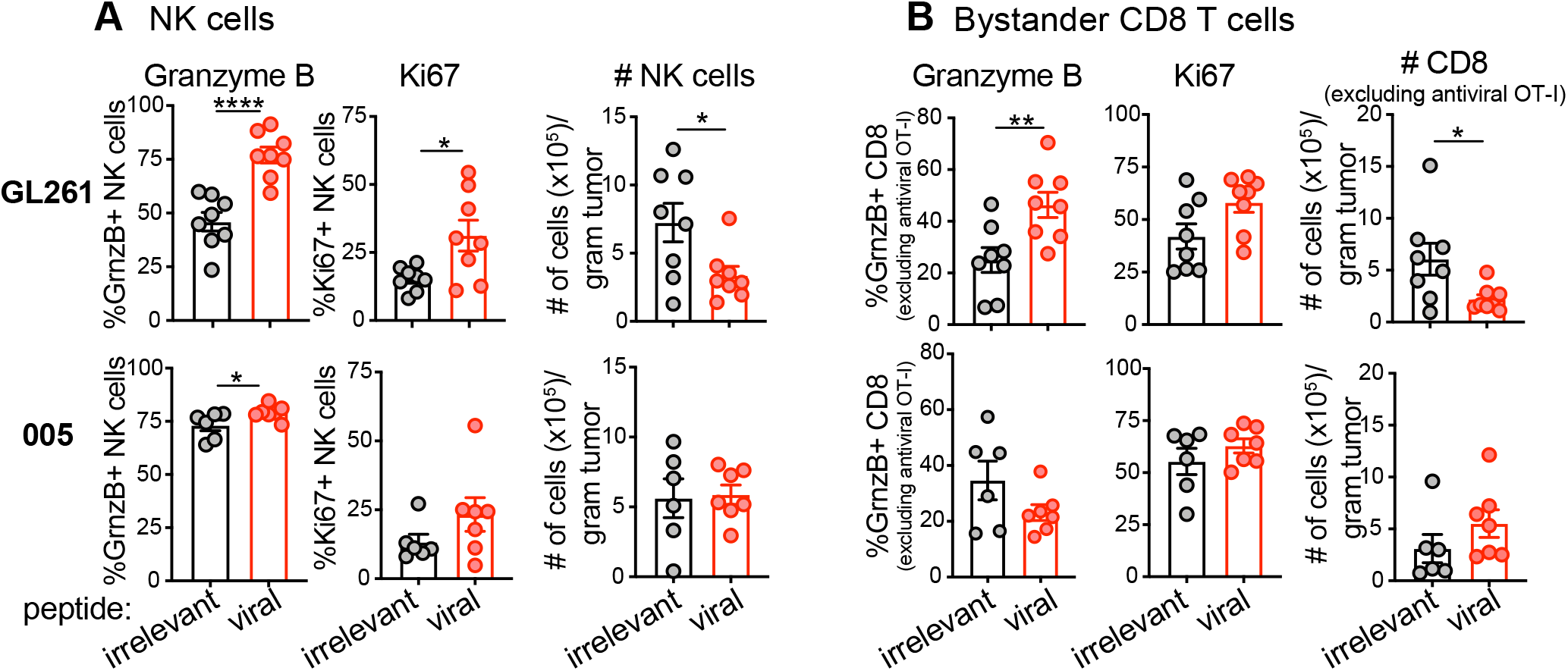
Virus-specific T_RM_ can perform sensing and alarm functions in mouse GBM. **A)** Proportion of NK cells expressing granzyme B and Ki67, and quantification of NK cells by flow cytometry in GL261 (top) or 005 (bottom) tumors 48h post-challenge with control or viral SIINFEKL peptide. **B)** Proportion of non-OT-I memory CD8+ T cells (gated on CD44+) expressing granzyme B and Ki67, and quantification of cells by flow cytometry in GL261 (top) or 005 (bottom) tumors 48h post-peptide challenge. Statistical significance was determined by unpaired two-tailed t-test (**A** and **B**, granzyme B) and unpaired two-tailed Mann-Whitney test (**B**, # CD8) where ^*^p<0.05, ^**^p<0.01, ^****^p<0.0001.

### Therapeutic efficacy of peptide alarm therapy

Given the observed immune activation following peptide alarm therapy, we hypothesized that antiviral T_RM_ reactivation could be leveraged as a GBM therapy to counteract the suppressive tumor environment and promote an anti-tumor response. To test this, we intracranially inoculated OT-I immune memory mice with GL261 or 005 tumors. We then performed peptide alarm therapy by intratumorally injecting viral-derived SIINFEKL peptide or irrelevant control peptide twice, 48hrs apart. We then assessed survival following therapy. We saw a significant increase in survival following treatment of mice with 005 tumors, but not GL261 (Figure 6). While all mice still succumbed to tumor, the median survival of mice bearing 005 tumors was extended by 5 days (32 to 37 days). These data contrast with the observed increase in immune activation in GL261 than in 005 tumors (Figs 4 and 5), indicating that, at least at the timepoints examined, the magnitude of CD8+ T or NK cell activation or accumulation in tumors is not necessarily predictive of therapeutic efficacy with PAT in GBM models. In all, these data show that peptide alarm therapy is effective at increasing survival of mice with GBM, but this may depend on the tumor model.

**Figure 6.**
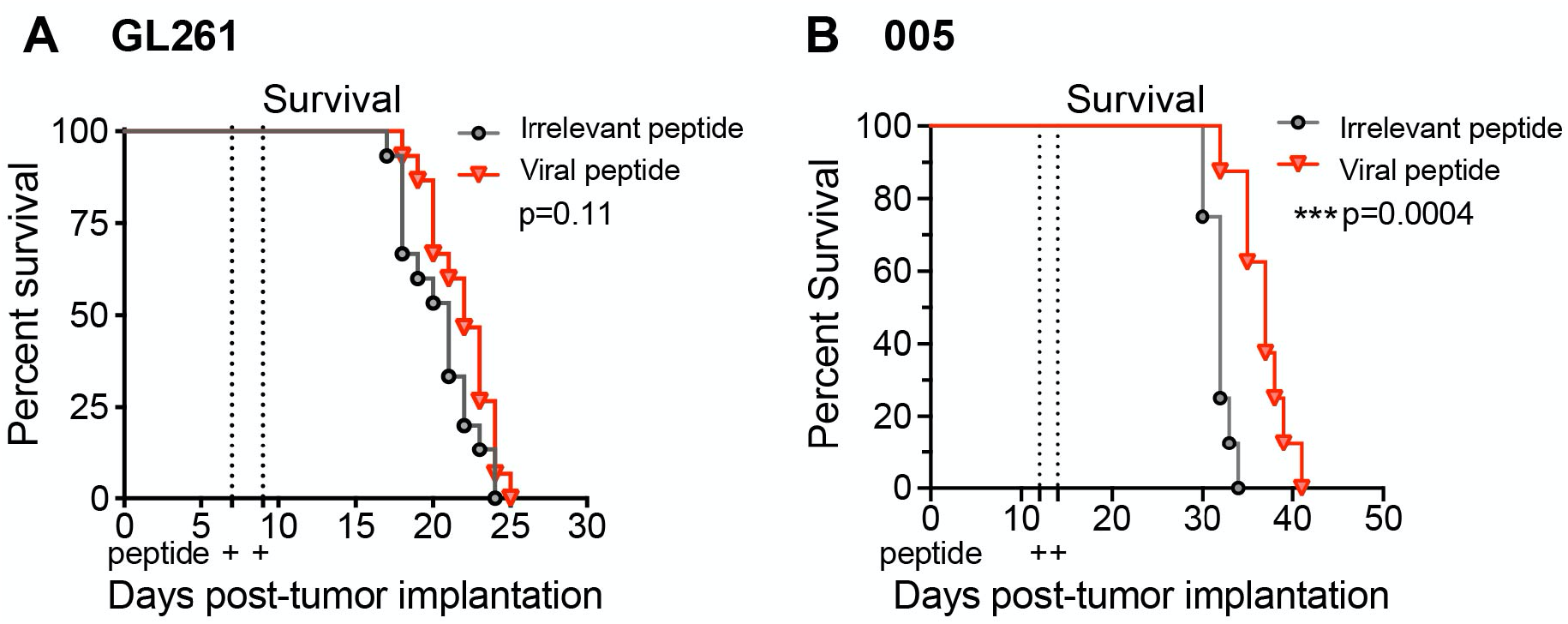
Therapeutic efficacy of peptide alarm therapy. Survival of mice with GL261 **(A)** or 005 **(B)** GBM tumors receiving two intratumoral injections of irrelevant (black) or viral SIINFEKL (red) peptides twice, 2 days apart. Statistical significance was determined by Log-rank Mantel-Cox test where ^***^p<0.001. **A)** n=15 mice pooled from 3 independent experiments. **B)** n=8 mice, pooled from 2 independent experiments.

## Discussion

This is the first study demonstrating an abundance of virus-specific T_RM_ in both mouse and human glioblastoma tumors, establishing these cells as a previously unappreciated but significant part of the GBM tumor immune environment. We further showed that these cells are functional in both human and mouse glioblastomas and that activating antiviral T_RM_ is a potential therapeutic approach.

Peptide alarm therapy (PAT) taps into the immune activating functions of T_RM_ by mimicking a local reinfection through delivery of viral-derived peptides in situ. Other groups have employed this strategy as a systemic therapy by conjugating viral-derived peptides to tumor-targeting antibodies, which has demonstrated efficacy against patient derived breast, liver and lung xenograft mouse tumor models (25,26). While this may be an attractive delivery approach in some cancers, systemic therapies are often restricted by the blood brain barrier and fail to effectively reach GBM. In a clinical setting, surgery is the standard of care for GBM patients, and it will be important to understand the translatability of PAT delivered to the resected tumor site at the time of surgery. Indeed, it’s been shown that CD8+ T cell responses change following surgical resection of glioblastoma tumors in mouse models (24).

While the role of CMV in GBM is controversial, several studies have found CMV antigens in human glioblastoma tumors (27). These findings have sparked clinical trials using dendritic cell vaccines to promote anti-CMV T cell responses (28), and CMV-specific T cell adoptive therapies (29,30). It is worth noting that we found no clear enrichment of CMV-specific T cells in GBM tumors, and moreover, identified EBV and IAV-specific T cells, suggesting broad localization of virus-specific T cells to GBM tumors.

It is interesting that despite an abundance of highly responsive antiviral T cells, we did not observe therapeutic efficacy in the GL261 model but did in 005 (Figure 6). Our GL261 results contrasts with other tumor models we have tested, including a subcutaneous MC38 colon carcinoma, intradermal B16 melanoma, and an autochthonous dermal melanoma model (3). This suggests that the magnitude of T_RM_ activation is not predicative of therapeutic efficacy, and that there may be glioblastoma or central nervous system (CNS) microenvironment influences that mitigate the downstream consequences of T_RM_ reactivation. A deeper understanding of these mechanisms will lend insights into poorly understood T_RM_ biology in the CNS as well as glioblastoma immuno-biology.

In healthy tissue and other murine tumor models, T_RM_ reactivation results in a significant increase in the number of virus-specific T cells, bystander CD8+ T cell and NK cells (3,12,31). Interestingly, this is also supported in humans by a recent study in hepatocellular carcinoma, which demonstrated a link between activated HBV-specific T_RM,_ and infiltration of bystander CD8+ T cells into the tumor (32). In contrast to these studies, we did not find an increase in T cells or NK cells in GBM tumors following T_RM_ activation, and even saw a *decrease* in some of these cell populations. This indicates that in GBM, canonical T_RM_ functions may not operate as they do in other tissues, and it may be that the restrictions imposed by the blood-brain barrier on immune cell recruitment are not overcome by T_RM_ reactivation in these mouse models. Understanding how immune cell homing to tumors following PAT occurs in other models (like melanoma) but not in GBM will be important for moving this therapy forward and will have implications for our fundamental understanding of T cell surveillance of the brain.

Investigating combination therapies that may synergize with PAT will be important in moving this therapy forward. For example, immune checkpoint inhibitors have poor demonstrated efficacy in GBM clinical trials (33), but we have previously shown that PAT sensitizes normally resistant murine B16 melanoma tumors resulting in tumor clearance (3). Oncolytic viruses (OV) also present an opportunity for possible therapeutic synergy; there are several active clinical trials testing OVs in glioblastoma, and many of these therapeutics are engineered from viruses to which the population has pre-existing immunity, such as herpes simplex virus (HSV), poliovirus, and measles virus (34). This presents an intriguing possibility that these therapies are reactivating pre-existing OV-specific T cells that were generated during prior infections. This also poses an opportunity to engineer OVs to express immunodominant epitopes from common viral infections to reactivate additional pre-existing memory T cells in tumors.

In summary, T_RM_ specific for common viral infections can reactivate in response to in situ delivered viral-derived peptides in murine glioblastoma models, and human glioblastoma explants. This reactivation occurred despite the immunosuppressive environment and was able to trigger NK cell immune activation and ultimately prolong survival of mice harboring 005 GBM tumors. This study supports the potential for leveraging virus-specific T_RM_ for GBM therapy.

## Acknowledgements

The authors thank Dr. Ryan Langlois for PR8-gp33, the NIH Tetramer Faculty for HLA-A^*^02:01 heavy chain plasmid, the Minnesota Supercomputing Institute for assistance with RNA sequencing, and University of Minnesota (UMN) BioNet for assistance with human samples. This work was supported by a UMN SPORE Program Project Planning grant (DM, CCC), NCI 1R01CA238439 (DM), Humor to Fight the Tumor Foundation (JN), and support from the Norris Cotton Cancer Center, NCI 5P30CA023108-42 (PR).

## Methods

### Mice

C57BL/6J (B6) female mice were purchased from The Jackson Laboratory (Bar Harbor, ME) and were maintained in specific-pathogen-free conditions at the University of Minnesota or Dartmouth College. CD90.1^+^ OT-I, CD45.1^+^ OT-I and Thy1.1+ P14 mice were fully backcrossed to C57BL/6J mice and maintained in our animal colony. Sample size was chosen on the basis of previous experience. No sample exclusion criteria were applied. No method of randomization was used during group allocation, and investigators were not blinded. All mice used in experiments were 5-14 weeks of age. All mice were used in accordance with the Institutional Animal Care and Use Committees guidelines at the University of Minnesota and Dartmouth College.

### Adoptive transfers and infections

Immune memory mice were generated by transferring 5×10^4^ CD90.1 or CD45.1 OT-I or CD90.1 P14 CD8^+^ T cells from female mice into naive 6-8 week old C57BL/6J female mice. One day following transfer, mice were infected with 1×10^6^ PFU of vesicular stomatitis virus expressing chicken ovalbumin (VSV_ova_) i.v., 5×10^4^ VSV_ova_ intranasally, 5×10^5^ PFU LCMV Armstrong intranasally, or 500 PFU of Influenza A strain PR8 expressing the gp33 epitope of LCMV (PR8_gp33_) intranasally.

### Lymphocyte isolation and phenotyping of mouse cells

We used an intravascular staining method to discriminate cells present in the vasculature from cells in the tissue parenchyma, as described for 12 hour timepoints (35). In brief, we injected mice i.v. with biotin/fluorochrome-conjugated anti-CD8α i.v. Three minutes after the injection, we euthanized the animals and harvested tissues as described^30^. GBM tumors were removed, digested in Collagenase IV (Sigma) with DNAse for 30 minutes then dissociated via gentleMACS Dissociator (Miltenyi Biotec) and lymphocytes purified on a 44/67% Percoll (GE Healthcare) gradient. Isolated mouse cells were stained with antibodies to CD103 (clone 2E7, eBioscience, 17-1031-80), NK1.1 (clone PK136, BioLegend, 108728), CD8a (clone 53-6.7, BioLegend, 100743), IFNγ (clone XMG1.2, BD Biosciences, 54411), CD25 (clone PC61, BD Biosciences, 557192), CD44 (clone IM7, BioLegend,103059), granzyme B (clone GB11, Invitrogen, GRB04), CD69 (clone H1.2F3, BD Biosciences, 562455), Ki67 (clone SolA15, Invitrogen, 25-5698-82). All cells were stained at antibody dilutions of 1:100 except for granzyme B (1:30). Cells stained intracellularly (for IFNγ, granzyme B and Ki67) were permeabilized using Tonbo or ebioscience Fixation/permeabilization kits. Cell viability was determined with Ghost Dye 780 (Tonbo Biosciences). Enumeration of cells was done using PKH26 reference microbeads (Sigma). The stained samples were acquired with LSRII or LSR Fortessa flow cytometers (BD) and analyzed with FlowJo software (Treestar).

### Immunofluorescence microscopy

GL261 or 005 tumors were harvested, then fixed in 2% paraformaldehyde for 2 h before being treated with 30% sucrose overnight for cryoprotection. The sucrose-treated tissue was embedded in OCT tissue-freezing medium and frozen in an isopentane liquid bath. Frozen blocks were processed, stained, and imaged including staining with antibodies to CD8-β (YTS156.7.7; BD Biosciences), CD90.1 (OX-7; BD Biosciences), and CD45.1 (A20; Biolegend). Sections were counterstained with 4′,6-diamidino-2-phenylindole dihydrochloride (DAPI) to detect nuclei.

### In situ tetramer staining

Human GBM tumors were sliced into thin sections manually with a sharp surgical blade. Sections were incubated overnight at 4 degrees C with 2μg/mL PE-conjugated HLA-A2 tetramer in PBS supplemented with 2% FBS and DNAse. Sections were then fixed in 2% paraformaldehyde for 2 hours before being treated with 30% sucrose overnight for cryoprotection. Tissues were then frozen and sectioned as described above. PE-tetramer was amplified with a polyclonal rabbit-anti-PE antibody at a 1:600 dilution (Novus Biologicals, NB120-7011) followed by goat-anti-rabbit Cy3 at a 1:800 dilution (Jackson ImmunoResearch, 111-165-003). Sections were also co-stained with antibodies to human CD8a and CD31, and counterstained with DAPI to detect nuclei.

### Tumor models and treatment

50,000-100,000 GL261 or 20,000 005 cells were injected intracranially into mice. Briefly, a burr hole, 1-1.5 mm in diameter was made using a hand drill without damaging the underlying dura mater. Cells were then injected using a Hamilton syringe guided by a stereotaxic frame into the cerebral hemisphere (3 mm deep, 1.8 mm to the right of bregma). Once the target was reached, 3ul of cells was slowly injected into the brain over a 5 minute period followed by 5 minutes rest, after which the syringe was slowly removed to avoid back-flow and the burr hole filled with bone wax. For survival studies, mice were monitored for clinical symptoms, and mice were euthanized when becoming moribund, and the presence of tumor was confirmed post-mortem. GL261 cells were maintained in DMEM (Life Technologies) supplemented with 10% Fetal Bovine Serum and penicillin/streptomycin (Cellgro), and 005 cells in DMEM/F12 medium (Life Technologies) with L-glutamine (2 mM; Corning), 1% N2 supplement (ThermoFisher Gibco), heparin (2 μg/mL; Sigma), penicillin/streptomycin (Cellgro), recombinant EGF (20 ng/mL; R&D Systems), and recombinant FGF2 (20 ng/mL;Peprotech).

For local tumor T cell reactivation experiments involving peptides, 0.5 µg of the indicated peptides (New England Peptides) were delivered by direct intratumor injection (as described above) in a volume of 3 µl phosphate buffered saline. Peptides used in mouse studies: KAVYNFATM (gp33) from LCMV (used as control/irrelevant) and SIINFEKL from ovalbumin.

### Procurement and processing of human blood and tissue samples

All tumor tissue and blood were obtained from male or female patients age 16-80 undergoing routine surgical resection of solid tumors or tumor metastases. Tumor tissue not required for pathological diagnostic procedures was obtained after surgical resection at the University of Minnesota and collected and de-identified by the Tissue Procurement Facility (BioNet, University of Minnesota). Informed consent was obtained from all subjects. The University of Minnesota Institutional Review Board approved all protocols used. Blood was collected in EDTA collection tubes and tumors were collected in RPMI media containing 5% FBS. All samples were stored at 4 degrees until processed (within 24 hours). Specimens reported on were obtained from HLA^*^A02+ patients that had sufficient tetramer+ cells for analysis by flow cytometry. Human blood was processed by Ficoll gradient. Tumors were minced and digested in Collagenase type IV with DNAse for 30 minutes then dissociated via gentleMACS Dissociator, and lymphocytes purified on a 44/67% Percoll (GE Healthcare) gradient. Lymphocytes were stained for anti-human HLA-A2 (clone BB7.2, BioLegend, 343324), CCR7 (clone G043H7, BioLegend, 353208), CD45RO (clone UCHL1, BioLegend, 304232), CD8α (clone SK1, BD Biosciences, 561945), CD3e (clone SP34-2, BD Biosciences, 557917), CD4 (clone L200, BD Biosciences, 551980), CD69 (clone FN50, BioLegend, 310926), CD103 (clone HML-1, Beckman Coulter, IM1856U). Cells were stained at antibody dilutions of 1:30. Samples were also stained for PE conjugated HLA-A^*^02 tetramers (made in house) for EBV_GLC_, EBV_CGL_, CMV_NLV_, and IAV_GIL_. Viability was assessed by live/dead staining with GhostDye510 (Tonbo biosciences).

### Transwell cultures and RNA isolation

Tumors were sliced into thin sections manually with a sharp surgical blade. Sections were then incubated in RPMI media containing 10% FBS, L-glutamine, sodium pyruvate, penicillin/streptomycin, HEPES, nonessential amino acids, and beta-mercapto-ethanol on 24-well polycarbonate Transwell inserts with a 0.4μm pore size (Corning) and maintained in 5% CO2 and atmospheric oxygen levels (15). Tissues were incubated with viral peptides at 10μg/mL or in equal volume of DMSO for 9 hours. Tissues with poor viability after culture were excluded. Tumor sections were stored in RNAlater (ThermoFisher) at 4 degrees overnight, then stored at -80 until further processing. For RNA isolation, tissue was thawed on ice in 1mL TRIZOL (Invitrogen) then homogenized with a Tissue Tearor homogenizer, BioSpec. RNA was then isolated following the TRIZOL recommended protocol. Resulting RNA was then further purified using Qiagen RNA Cleanup Kit. Peptides used in human studies: CLGGLLTMV (EBV_CLG_), GLCTLVAML (EBV_GLC_), NLVPMVATV (CMV_NLV_), GILGFVFTL (IAV_GIL_).

### RNA library preparation and sequencing

mRNA libraries were generated using the TruSeq Stranded mRNA Library Prep kit (Illumina) and sequenced on an Illumina HiSeq 2500 in 50-base paired-end reactions. Fastq files were verified for quality control using the fastqc software package. Low-quality segments and adapters were trimmed using Trimmomatic. Quality-filtered reads were aligned to the human genome GRCh38 using Hisat (36). Differentially expressed genes were determined using the DESeq2 R package (37) where false-discovery rate (FDR) <0.05 was considered significant. Upstream transcriptional regulators, canonical pathways and summary plots were generated through the use of Ingenuity Pathway Analysis (IPA, QIAGEN Inc., https://www.qiagenbioinformatics.com/products/ingenuity-pathway-analysis) (38).

### Statistics

Data were subjected to the Shapiro-Wilk normality test to determine whether they were sampled from a Gaussian distribution. If a Gaussian model of sampling was satisfied, parametric tests (unpaired two-tailed Student’s *t*-test for two groups and one-way ANOVA with Bonferroni multiple comparison test for more than two groups) were used. If the samples deviated from a Gaussian distribution, non-parametric tests (Mann–Whitney U test for two groups, Kruskal–Wallis with Dunn’s multiple comparison test for more than two groups). Survival data were analyzed by Kaplan Meier survival curves, and comparisons determined by log-rank (Mantel-Cox) test. All statistical analysis was done in GraphPad Prism (GraphPad Software Inc.). *P* < 0.05 was considered significant.

### Cell definitions

T_RM_ : In this study, we have defined T_RM_ as virus-specific memory CD8+ T cells, identified by tetramer or use of TCR transgenic T cells, expressing CD69.

### Data availability

RNAseq data available upon request.

## References

1. Yu MW, Quail DF. Immunotherapy for Glioblastoma: Current Progress and Challenge. Front Immunol. 2021;12:676301.

2. Lim M, Xia Y, Bettegowda C, Weller M. Current state of immunotherapy for glioblastoma. Nature Reviews Clinical Oncology. 2018;15:422–42.

3. Rosato PC, Wijeyesinghe S, Stolley JM, Nelson CE, Davis RL, Manlove LS, et al. Virus-specific memory T cells populate tumors and can be repurposed for tumor immunotherapy. Nat Commun. Nature Publishing Group; 2019;10:567.

4. Scheper W, Kelderman S, Fanchi LF, Linnemann C, Bendle G, MAJ de Rooij, et al. Low and variable tumor reactivity of the intratumoral TCR repertoire in human cancers. Nature Medicine. 2018;359:1350–94.

5. Chiou S-H, Tseng D, Reuben A, Mallajosyula V, Molina IS, Conley S, et al. Global analysis of shared T cell specificities in human non-small cell lung cancer enables HLA inference and antigen discovery. Immunity. 2021;54:586-602.e8.

6. Duhen T, Duhen R, Montler R, Moses J, Moudgil T, de Miranda NF, et al. Co-expression of CD39 and CD103 identifies tumor-reactive CD8 T cells in human solid tumors. Nature Communications. 2018;9:56.

7. Andersen RS, Thrue CA, Junker N, Lyngaa R, Donia M, Ellebæk E, et al. Dissection of T-cell antigen specificity in human melanoma. Cancer Res. 2012;72:1642–50.

8. Simoni Y, Becht E, Fehlings M, Loh CY, Koo S-L, Teng KWW, et al. Bystander CD8 + T cells are abundant and phenotypically distinct in human tumour infiltrates. Nature. 2018;557:575.

9. Schenkel JM, Fraser KA, Vezys V, Masopust D. Sensing and alarm function of resident memory CD8^+^ T cells. Nat Immunol. 2013;14:509–13.

10. Rosato PC, Beura LK, Masopust D. Tissue resident memory T cells and viral immunity. Current Opinion in Virology. 2017;22:44–50.

11. Rosato PC, Wijeyesinghe S, Stolley JM, Masopust D. Integrating resident memory into T cell differentiation models. Current Opinion in Immunology. 2020;63:35–42.

12. Schenkel JM, Fraser KA, Beura LK, Pauken KE, Vezys V, Masopust D. Resident memory CD8 T cells trigger protective innate and adaptive immune responses. Science. 2014;346:98–101.

13. Ariotti S, Hogenbirk MA, Dijkgraaf FE, Visser LL, Hoekstra ME, Song J-Y, et al. T cell memory. Skin-resident memory CD8^+^ T cells trigger a state of tissue-wide pathogen alert. Science. 2014;346:101–5.

14. Szabo PA, Miron M, Farber DL. Location, location, location: Tissue resident memory T cells in mice and humans. Sci Immunol. 2019;4:eaas9673.

15. Davies EJ, Dong M, Gutekunst M, Närhi K, HJAA van Zoggel, Blom S, et al. Capturing complex tumour biology in vitro: histological and molecular characterisation of precision cut slices. Scientific Reports. 2015;5:17187.

16. Beura LK, Hamilton SE, Bi K, Schenkel JM, Odumade OA, Casey KA, et al. Normalizing the environment recapitulates adult human immune traits in laboratory mice. Nature. 2016;532:512–6.

17. Wakim LM, Woodward-Davis A, Liu R, Hu Y, Villadangos J, Smyth G, et al. The Molecular Signature of Tissue Resident Memory CD8 T Cells Isolated from the Brain. The Journal of Immunology. 2012;189:1201305–3471.

18. Wijeyesinghe S, Beura LK, Pierson MJ, Stolley JM, Adam OA, Ruscher R, et al. Expansible residence decentralizes immune homeostasis. Nature. 2021;

19. Hawke S, Stevenson PG, Freeman S, Bangham CR. Long-term persistence of activated cytotoxic T lymphocytes after viral infection of the central nervous system. The Journal of Experimental Medicine. 1998;187:1575–82.

20. Urban SL, Jensen IJ, Shan Q, Pewe LL, Xue H-H, Badovinac VP, et al. Peripherally induced brain tissue–resident memory CD8 + T cells mediate protection against CNS infection. Nature Immunology. Nature Publishing Group; 2020;21:938–49.

21. Nelson CE, Thompson EA, Quarnstrom CF, Fraser KA, Seelig DM, Bhela S, et al. Robust Iterative Stimulation with Self-Antigens Overcomes CD8+ T Cell Tolerance to Self- and Tumor Antigens. Cell Reports. 2019;28:3092-3104.e5.

22. Szatmári T, Lumniczky K, Désaknai S, Trajcevski S, Hídvégi EJ, Hamada H, et al. Detailed characterization of the mouse glioma 261 tumor model for experimental glioblastoma therapy. Cancer Science. 2006;97:546–53.

23. Marumoto T, Tashiro A, Friedmann-Morvinski D, Scadeng M, Soda Y, Gage FH, et al. Development of a novel mouse glioma model using lentiviral vectors. Nature Medicine. Nature Publishing Group; 2009;15:110–6.

24. Khalsa JK, Cheng N, Keegan J, Chaudry A, Driver J, Bi WL, et al. Immune phenotyping of diverse syngeneic murine brain tumors identifies immunologically distinct types. Nature Communications. Nature Publishing Group; 2020;11:3912.

25. Millar DG, Ramjiawan RR, Kawaguchi K, Gupta N, Chen J, Zhang S, et al. Antibody-mediated delivery of viral epitopes to tumors harnesses CMV-specific T cells for cancer therapy. Nature Biotechnology. Nature Publishing Group; 2020;38:420–5.

26. Sefrin JP, Hillringhaus L, Mundigl O, Mann K, Ziegler-Landesberger D, Seul H, et al. Sensitization of Tumors for Attack by Virus-Specific CD8+ T-Cells Through Antibody-Mediated Delivery of Immunogenic T-Cell Epitopes. Front Immunol. 2019;10:1962.

27. Rahman M, Dastmalchi F, Karachi A, Mitchell D. The role of CMV in glioblastoma and implications for immunotherapeutic strategies. Oncoimmunology. 2018;8:e1514921.

28. Batich KA, Mitchell DA, Healy P, Herndon JE, Sampson JH. Once, Twice, Three Times a Finding: Reproducibility of Dendritic Cell Vaccine Trials Targeting Cytomegalovirus in Glioblastoma. Clin Cancer Res. American Association for Cancer Research; 2020;26:5297–303.

29. Smith C, Lineburg KE, Martins JP, Ambalathingal GR, Neller MA, Morrison B, et al. Autologous CMV-specific T cells are a safe adjuvant immunotherapy for primary glioblastoma multiforme. J Clin Invest. 2020;130:6041–53.

30. Weathers S-P, Penas-Prado M, Pei B-L, Ling X, Kassab C, Banerjee P, et al. Glioblastoma-mediated Immune Dysfunction Limits CMV-specific T Cells and Therapeutic Responses: Results from a Phase I/II Trial. Clin Cancer Res. American Association for Cancer Research; 2020;26:3565–77.

31. Beura LK, Mitchell JS, Thompson EA, Schenkel JM, Mohammed J, Wijeyesinghe S, et al. Intravital mucosal imaging of CD8+resident memory T cells shows tissue-autonomous recall responses that amplify secondary memory. Nature Immunology. 2018;19:173–82.

32. Cheng Y, Gunasegaran B, Singh HD, Dutertre C-A, Loh CY, Lim JQ, et al. Non-terminally exhausted tumor-resident memory HBV-specific T cell responses correlate with relapse-free survival in hepatocellular carcinoma. Immunity. 2021;54:1825-1840.e7.

33. Akintola OO, Reardon DA. The Current Landscape of Immune Checkpoint Blockade in Glioblastoma. Neurosurg Clin N Am. 2021;32:235–48.

34. Liu P, Wang Y, Wang Y, Kong Z, Chen W, Li J, et al. Effects of oncolytic viruses and viral vectors on immunity in glioblastoma. Gene Ther. 2020;1–12.

35. Anderson KG, Mayer-Barber K, Sung H, Beura L, James BR, Taylor JJ, et al. Intravascular staining for discrimination of vascular and tissue leukocytes. Nat Protoc. 2014;9:209–22.

36. Kim D, Langmead B, Salzberg SL. HISAT: a fast spliced aligner with low memory requirements. Nature methods. 2015;12:357–60.

37. Love MI, Huber W, Anders S. Moderated estimation of fold change and dispersion for RNA-seq data with DESeq2. Genome Biology. 2014;15:550.

38. Krämer A, Green J, Pollard J, Tugendreich S. Causal analysis approaches in Ingenuity Pathway Analysis. Bioinformatics. 2014;30:523–30.

